# Stability of local secondary structure determines selectivity of viral RNA chaperones

**DOI:** 10.1101/293191

**Authors:** Jack P.K. Bravo, Alexander Borodavka, Anders Barth, Antonio N. Calabrese, Peter Mojzes, Joseph J.B. Cockburn, Don C. Lamb, Roman Tuma

## Abstract

To maintain genome integrity, segmented double-stranded RNA viruses of the *Reoviridae* family must accurately select and package a complete set of up to a dozen distinct genomic RNAs. It is thought that the high fidelity segmented genome assembly involves multiple sequence-specific RNA-RNA interactions between single-stranded RNA segment precursors. These are mediated by virus-encoded non-structural proteins with RNA chaperone-like activities, such as rotavirus NSP2 and avian reovirus σNS. Here, we compared the abilities of NSP2 and σNS to mediate sequence-specific interactions between rotavirus genomic segment precursors. Despite their similar activities, NSP2 successfully promotes inter-segment association, while σNS fails to do so. To understand the mechanisms underlying such selectivity in promoting inter-molecular duplex formation, we compared RNA-binding and helix-unwinding activities of both proteins. We demonstrate that octameric NSP2 binds structured RNAs with high affinity, resulting in efficient intramolecular RNA helix disruption. Hexameric σNS oligomerises into an octamer that binds two RNAs, yet it exhibits only limited RNA-unwinding activity compared to NSP2. Thus, the formation of intersegment RNA-RNA interactions is governed by both helix-unwinding capacity of the chaperones and stability of RNA structure. We propose that this protein-mediated RNA selection mechanism may underpin the high fidelity assembly of multi-segmented RNA genomes in *Reoviridae*.

## Introduction

Members of the *Reoviridae* family of double-stranded (ds)RNA viruses encompass important human, animal and plant pathogens (1). Reovirus genomes are distributed between 9–12 RNA segments, all of which are essential for virus assembly and replication (2). Multiple sequence-specific RNA-RNA interactions between ssRNA segment precursors are believed to underpin the assembly of a complete multi-segmented viral genome (3–7). The underlying molecular mechanisms of high fidelity RNA selection and accurate genome assembly remain poorly understood (1, 3). Segment selection, genome replication and virus assembly occur in cytoplasmic membraneless organelles, termed viral factories or viroplasms (8–11). Non-structural proteins rotavirus (RV) NSP2, or mammalian and avian reovirus (ARV) σNS are major components of viroplasms and viral factories, and are essential for genome replication and virus assembly (8, 12–16).

Recently, we have demonstrated that NSP2 can promote selective RNA duplex formation between genomic ssRNAs, acting as an RNA chaperone (17). We have shown previously that ARV σNS exhibits similar non-specific ssRNA binding, helix-destabilizing and strand-annealing activities to NSP2, suggesting that both proteins may play similar roles during genome assembly (18). Despite these apparent similarities, it remains unclear whether these proteins employ similar mechanisms of facilitating specific inter-segment interactions.

Here, using a previously established interaction between RV segments S5 and S11 ssRNAs as a model system, we compared the abilities of NSP2 and σNS to promote intersegment RNA duplex formation. This allowed us to decouple the RNA chaperone activities of both proteins from their multiple, overlapping roles during virus replication, such as viroplasm assembly, kinase activity of NSP2 and interactions with other viral and cellular proteins (8, 12–16). While NSP2 is efficient at mediating stable S5: S11 RNA-RNA interactions, σNS completely fails to do so. While both proteins simultaneously bind multiple unstructured ssRNAs with similar affinities, NSP2 has a greater propensity for binding and unfolding structured RNA hairpins. RNA stem-loop unwinding experiments monitored by single-pair FRET indicates that NSP2 disrupts RNA secondary structure more efficiently, whereas σNS binding induces an ensemble of partially unfolded intermediates. Upon binding multiple RNAs, hexameric σNS undergoes further oligomerisation, forming stable octameric ribonucleoprotein (RNP) complexes with increased helix-destabilizing activity. Such differences in modes of RNA binding and helix-unwinding between these proteins may explain the failure of σNS to promote a stable inter-segment S5: S11 RNA-RNA interaction. These results suggest that RNA structural stability can modulate viral RNA chaperone activity, thereby restricting non-cognate RNA duplex formation during segment selection and genome assembly.

## Materials and Methods

### Protein expression and purification

NSP2 and σΝS were expressed and purified as previously described in (18). Size exclusion chromatography (SEC) analysis was performed using Superdex S200 10/300 column (GE healthcare) equilibrated in SEC buffer (50mM HEPES pH 8.0, 150 mM NaCl) at 4°C.

### RNA production

RV segments S5 and S11 were produced and labelled as previously described in *Borodavka et al*, 2016 (19). AlexaFluor488 dye-labelled 20mer and unlabelled 15mer, 20mer and 40mer RNAs (**Supporting Table S1**) were purchased from Integrated DNA technologies (IDT). Cy3-and Cy5-labelled 17mer RNAs (**Supporting Table S1**) derived from the rotavirus segment S11 (nucleotides 49 – 66, complementary to segment S5 nucleotides 307 – 324) were ordered from IDT. Dual-labelled RNA stem-loop was designed to have a similar minimal free energy (MFE) of folding to that of the stable 20mer hairpin (**Supporting Table S1**), but with a larger loop to enable efficient protein binding. A 36mer hairpin with MFE = −8.9 kcal mol^-1^ with 3′-Atto532 and 5′-Atto647N dyes was designed and purchased from IBA Life Sciences.

### Size-exclusion chromatography (SEC), dynamic light scattering (DLS) and small angle X-ray scattering (SAXS)

σNS-ribonucleoprotein (RNP) complex was prepared by mixing σNS (175 µM monomer) with 20mer unstructured RNA (70 µM, **Supporting Table S1**) to ensure complete saturation of σNS with RNA, assuming that a single σNS hexamer binds two RNA molecules. In the case of a higher-order σNS-RNP complex (i.e. octamer bound to 2 RNAs), this stoichiometry also ensures protein saturation with RNA. For light scattering analysis, 25 µM σNS and σNS-20mer complex were run at a flow-rate of 0.4 ml min^-1^ on a TSKgel G6000PWxl SEC column (Tosoh) with an AKTA pure system (GE Healthcare) connected to a DAWN HELEOS (Wyatt).

SAXS samples were prepared in a similar manner, whereby σNS was incubated with saturating amounts of 20mer or 40mer RNA **(Supporting Table S1**), as described above. SAXS intensity data, *I(q)* versus momentum transfer *q* (*q* = 4*π*sin*θ/λ*, where *θ* is the scattering angle and λ is the wavelength), were collected using SEC-SAXS on beamline B21 at Diamond Light Source (Didcot, UK) over a range of *q* of 0.004 < *q* < 0.442 Å^-1^. 50 µl of each sample (~175 µM) was loaded onto a 2.4 ml Superdex 200 Increase 3.2 column mounted on Agilent HPLC, and the eluent was flowed through the SAXS beam at 0.04 ml/min. The SEC buffer used as the background was collected after one SEC column volume. SAXS data were collected at 1 second intervals using a Pilatus 2M detector (Dectris, Switzerland) at a distance of 3.9 m and an X-ray wavelength of 1 Å. Guinier plot fit and real space inversions were performed using Primus (20) and GNOM from the ATSAS software package v. 2.8.3 (21, 22). Radii of gyration (*R_g_*) were estimated using AUTORG (21). Low resolution envelopes were determined using the simulated annealing procedures implemented in DAMMIF (23) in slow mode, with no symmetry applied. Each scattering curve generated 20 independent models, which were averaged and filtered using DAMAVER (24) and DAMFILT with a mean normalized spatial discrepancy of 0.820 ± 0.05 (σNS apoprotein) and 0.576 ± 0.03 (σNS-20mer complex). SAXS experimental data together with the relevant experimental conditions and the derived models are available from SASBDB.

### Native mass spectrometry

σNS was dialysed into 200 mM ammonium acetate, pH 7.6 overnight at 4°C. RNP complexes were assembled as described above. σNS apoprotein and σNS-RNP complex were diluted to a final concentration of 20 µM. NanoESI–IMS–MS spectra were acquired with a Synapt HDMS mass spectrometer (Waters) with platinum/gold-plated borosilicate capillaries prepared in-house. Typical instrument parameters were: capillary voltage, 1.2–1.6 kV; cone voltage, 40 V; trap collision voltage, 6 V; transfer collision voltage, 10 V; trap DC bias, 20 V; backing pressure, 4.5 mbar; IMS gas pressure, 0.5 mbar; traveling wave height, 7 V; and traveling wave velocity, 250 ms^-1^. Data were processed with MassLynx v4.1, Driftscope 2.5 (Waters) and Massign (25). Collisional Cross-Sections (CCSs) were estimated through a calibration (26–28) using arrival-time data for ions with known CCSs (β-lactoglobulin A, avidin, concanavilin A and yeast alcohol dehydrogenase, all from Sigma Aldrich). The CCS values of the lowest observed charge state (and therefore the least affected by Coulombic repulsion (29)) were selected for comparison with SAXS-derived CCS estimates. Theoretical CCS values for SAXS *ab initio* reconstructions of σNS and σNS-RNP were generated by using the calibrated projection approximation method in IMPACT (30). Estimated CCSs for the 20 independently generated dummy atom models were generated and averaged.

### Affinity measurements by fluorescence anisotropy

Fluorescence anisotropy measurements with AlexaFluor488 dye-labelled RNAs (**Supporting Table S1**) were performed at 25ºC using a POLARstar Omega plate reader (BMG Labtech) in Greiner 384 well black polypropylene plates. Serial 2-fold dilutions of NSP2 and σNS were titrated into 5 nM RNA in 50 mM Tris-HCl pH 7.5, 50 mM NaCl, 1 mM EDTA, 0.05% Tween-20 in a total volume of 50 µl and equilibrated at room temperature for 15 minutes prior to measurements were taken. Where required, buffers were supplemented with 10 mM MgCl_2_. Raw Anisotropy (r) values were calculated as follows:

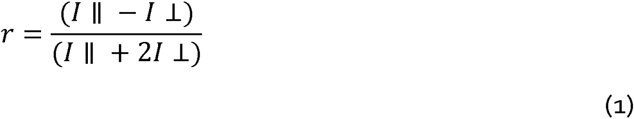

(1) Where *I ║* and *I* ⊥ are the parallel and perpendicular emission signals, respectively. Normalized anisotropy values were plotted as a function of protein concentration and fitted to a Hill equation using OriginPro 9.0.

### Electrostatic contributions to free energies of protein-RNA interactions

The dependence of K_obs_ on buffer ionic strength can be expressed as

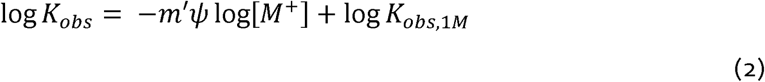

where [*M*^+^] is the monovalent counterion concentration (in this case Na^+^), *m′* is the number of ion pairs formed, *ψ* is defined as the thermodynamic extent of counterion binding and *K_obs,1M_* is the non-electrostatic contribution to the dissociation constant, defined as the *K_obs_* at 1 M NaCl, when the polyelectrolyte effect is minimal (31, 32). The slope, SK_obs_, of log(*K*_obs_) against log[*M*^+^] relates to the number of counterions released upon binding as follows:

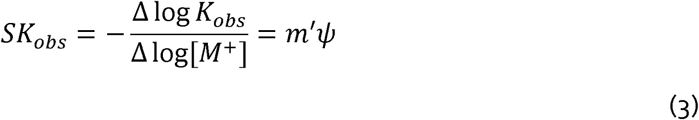

The electrostatic (poly-electrolyte) contribution to free energy of binding, ∆G_PE_, can therefore be determined as

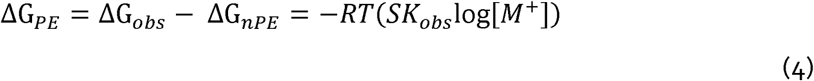

where ∆G_obs_ is the total free energy of binding, and ∆G_nPE_ is the non-electrostatic (non-polyelectrolyte) contribution. As the parameter *ψ* varies between different polynucleotides, we estimated the amount of salt bridges using empirically determined values, poly(U) = 0.68 and poly(A) = 0.78 as the upper and lower limits of salt bridges involved in complex formation (31, 32). As NSP2 octamers disassemble under high ionic strengths (33), a range of 50 – 250 mM NaCl was used to determine RNA binding affinities. σNS, however, aggregates at low (<50 mM) NaCl concentrations, but is not affected by higher ionic strength conditions, so a wider range of ionic strength buffers were used to determine RNA binding affinities (up to 500 mM NaCl).

### Electrophoretic Mobility Shift Assays (EMSAs) and in-gel FRET

A dual-labelled RNA stem-loop was designed with a minimum free energy (MFE) of folding of −8.9 kcal mol^-1^, containing 3′-donor (ATTO532) and 5′-acceptor (ATTO647N) dye fluorophores as a FRET pair (**Supporting Table S1**). Dual-labelled stem-loop was heat-annealed in binding buffer (50 mM Tris-HCl pH 7.5, 50 mM NaCl, 1 mM EDTA) at 75°C for 5 minutes and cooled to 4°C prior to incubation with increasing amounts of σNS (final RNA concentration 10 nM). 10 µl of each sample was mixed with 2 µl 6x loading buffer (30% v/v glycerol, 5% Ficoll 400, 50 mM NaCl, 10 mM HEPES pH 8, 5 mM EDTA, 0.002% w/v bromophenol blue). Electrophoresis was carried out on a nondenaturing 1.5% agarose gel for 90 min at 100 V in 1 X Tris-borate-EDTA buffer at 4°C. Gels were imaged using a fluorescence scanner (Fujifilm FLA-5100) with 532 nm excitation, imaging donor and acceptor wavelengths separately. 2D densitometry was performed using ImageJ. Apparent FRET efficiencies (E*_FRET(app)_*) were calculated as follows:

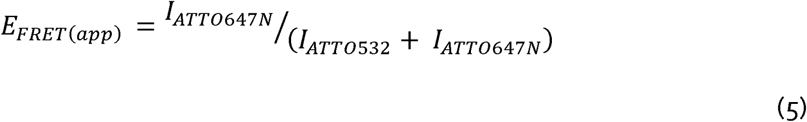

Where *I_ATTO532_* and *I_ATTO647N_* are donor and acceptor emission intensities, respectively. For probing RNA: RNA interactions between S5f and S11f fragments (**Supporting Table S1**), 100 nM of each RNA was individually heat-annealed at 85ºC for 5 minutes in folding buffer (10 mM HEPES pH 7, 1M NaCl, 10 mM MgCl_2_) and snap-cooled to 4ºC for 20 minutes, ensuring complete stem-loop folding prior to co-incubation. 10 nM of S5f and S11f were incubated at 37ºC in annealing buffer (10 mM HEPES pH 7, 100 mM NaCl, 1 mM MgCl_2_) and immediately analysed by electrophoresis on a native 15% acrylamide-TBE gel, run at 80 V at 4ºC, and stained with 0.01% (w/v) SYBR gold. Gels were imaged using a fluorescence scanner as described above, using 488 nm excitation.

### Fluorescence cross-correlation spectroscopy (FCCS)

Equimolar amounts of 18-nt Cy3-and Cy5-labelled non-complementary RNAs (10 nM each) (**Supporting Table S1**) were incubated with varying concentrations of NSP2 and σNS in 50 mM NaCl, 20 mM HEPES pH 7.4. Interactions between S5 and S11 RNAs and S5 and Cy5–17mer **(Supporting Table S1**) were measured as previously described (17). Briefly, 55 nM of each RNA strand was incubated with 5 – 10 µM NSP2 or σNS at 37ºC for 15 minutes. Samples were then diluted into 150 mM NaCl, 20 mM HEPES pH 7.4, 0.05% Tween 20, resulting in a final RNA concentration of 1 nM each labelled RNA. NSP2 removal using proteinase K does not significantly reduce the amplitude of cross-correlation, suggesting that the observed cross-correlation is due to strand-annealing (17).

### Circular Dichroism (CD)

CD experiments were performed on a Chirascan plus spectrometer (Applied Photophysics). Samples were prepared by dialyzing protein solutions against 10 mM phosphate buffer pH 7.4, 50 mM sodium fluoride. Spectra were recorded over a wavelength range of 190 – 260 nm, with a bandwidth of 1 nm, step size of 1 nm and a path length of 1 mm. An average of 3 scans were used for the final spectra. Thermal stability was analysed by monitoring the CD signal at 222 nm during heating from 20°C to 70°C with a heating rate of 1°C min^-1^.

### Ensemble FRET

Dual-labelled stem-loop (**Supporting Table S1**) was heat-annealed at 75°C for 5 minutes and cooled to 4°C. Folded RNA-alone and denatured RNA (in 50% v/v formamide) were initially measured in 100 µl volumes at a final RNA concentration of 10 nM. Serial 2-fold dilutions of NSP2 and σNS from 15 µM were incubated with 10 nM RNA at room temperature for 15 minutes prior to measurement. Measurements were performed using a Fluorolog spectrofluorimeter (Horiba Jobin-Yvon). Apparent FRET efficiencies were calculated using **Equation 5**.

### Single-pair FRET measurements

Dual-labelled RNA stem-loop used for ensemble FRET measurements was heat-annealed as described above and diluted to 10 pM final concentration. σNS and NSP2 were incubated with the stem-loop (25 nM each). Single-pair FRET measurements were performed on a custom-built confocal microscope with multiparameter fluorescence detection (MFD) and pulsed interleaved excitation (PIE) (34) as described previously (35). Briefly, two picosecond pulsed lasers (532 nm and 640 nm) were operated at a repetition rate of 26.66 MHz and delayed by 18 ns with respect to each other to achieve rapid alternating excitation of donor and acceptor fluorophore at 100 µW laser power. By diluting the sample to picomolar concentrations, single molecule events were detected as bursts of fluorescence as they diffuse through the confocal volume on the millisecond timescale. Bursts were selected using a sliding time window burst search (36) with a count rate threshold of 10 kHz, a time window of 500 µs and a minimum of 100 photons per burst. Using time-correlated single photon counting and polarized detection, one can calculate for every molecule its FRET efficiency, labeling stoichiometry, and the fluorescence lifetime and anisotropy of the donor and acceptor fluorophores (34). To remove molecules lacking the donor or acceptor dye, we used the ALEX-2CDE filter (37) with a time constant of 100 µs and an upper threshold of 10. Accurate FRET efficiencies *E* were calculated from background-corrected photon counts in the donor channel and acceptor channel after donor excitation (*FDD/FDA*) and acceptor channel after acceptor excitation (*F AA*) by correcting for crosstalK(*α=*0.03), direct excitation (*δ=*0.06) and differences in the quantum yield and detection efficiency (γ*=*0.65):

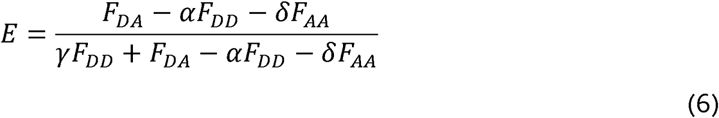

Species-selective fluorescence correlation functions were determined as follows: Sub-populations of molecules were selected using FRET efficiency thresholds (NSP2: low-FRET *E* < 0.4, high-FRET *E*> 0.6; σNS: low-FRET *E* < 0.15, medium-FRET 0.3 < *E* < 0.8, high-FRET *E*> 0.9). For every burst, photons in a time window of 50 ms around the edges of the burst were added. If another single molecule event was found in the time window, the respective burst is excluded from the analysis. Correlation functions were calculated for each individual burst using acceptor photons after acceptor excitation to ensure that the obtained correlation functions are independent of the FRET efficiency. Species-selective correlation functions were then averaged to obtain the burst-selective correlation function (38, 39). All analysis was performed using the *PAM* software package (40).

### Raman spectroscopy

Raman spectra of RNA, σNS and RNP and their corresponding buffers were acquired on a modular multi-channel Raman spectrograph Jobin Yvon–Spex 270M in 90° scattering geometry using 532 nm line of a continuous-wave solid-state Nd:YVO_4_ laser for excitation (power of 240 mW at the sample), as described in detail elsewhere (41). Raman measurements were performed in a temperature-controlled hermetical quartz microcell (4 µL volume) at 20°C and 60°C. Final spectra represent averages of 30 −720 individually acquired and analysed scans (depending on the sample type) each of 1 min integration time to notice any spectral changes during laser exposure and to increase signal-to-noise ratio without mathematical smoothing. Wavenumber scales were precisely calibrated (± 0.1 cm^-1^) using the emission spectra of a neon glow lamp taken before and after each Raman measurement. The Raman contribution from corresponding buffer was subtracted, and the spectra were corrected for non-Raman background.

## Results

### σNS is unable to promote inter-segment interactions between RV RNAs

Recently, we have demonstrated that NSP2 can selectively promote RNA-RNA duplex formation between genomic ssRNAs in rotaviruses (17). Both NSP2 and σNS possess helix-destabilising and strand-annealing activities *in vitro* (17, 18). Using a previously established RNA-RNA interaction between rotavirus (RV) segment S5 and S11 ssRNAs (17), we compared the abilities of NSP2 and σNS to promote inter-segment duplex formation. We employed fluorescence cross-correlation spectroscopy (FCCS) to monitor inter-segment RNA-RNA interactions.

While S5 and S11 do not interact in the absence of protein (**Figure 1A**, magenta), co-incubation of S5 and S11 with NSP2 results in inter-molecular RNA duplex formation (**Figure 1A**, blue). In contrast, co-incubation of S5 and S11 RNAs with σNS does not promote duplex formation (**Figure 1A**, black). Binding of both NSP2 and σNS to either RNA results in an increase in apparent diffusion time, confirming that both proteins interact with S5 and S11 and form larger ribonucleoprotein (RNP) complexes (**Supplementary Figure S1**). Hence, both proteins can bind to these RNAs, but only NSP2 can mediate a sequence-specific interaction between RV genomic ssRNAs.

**Figure. 1.**
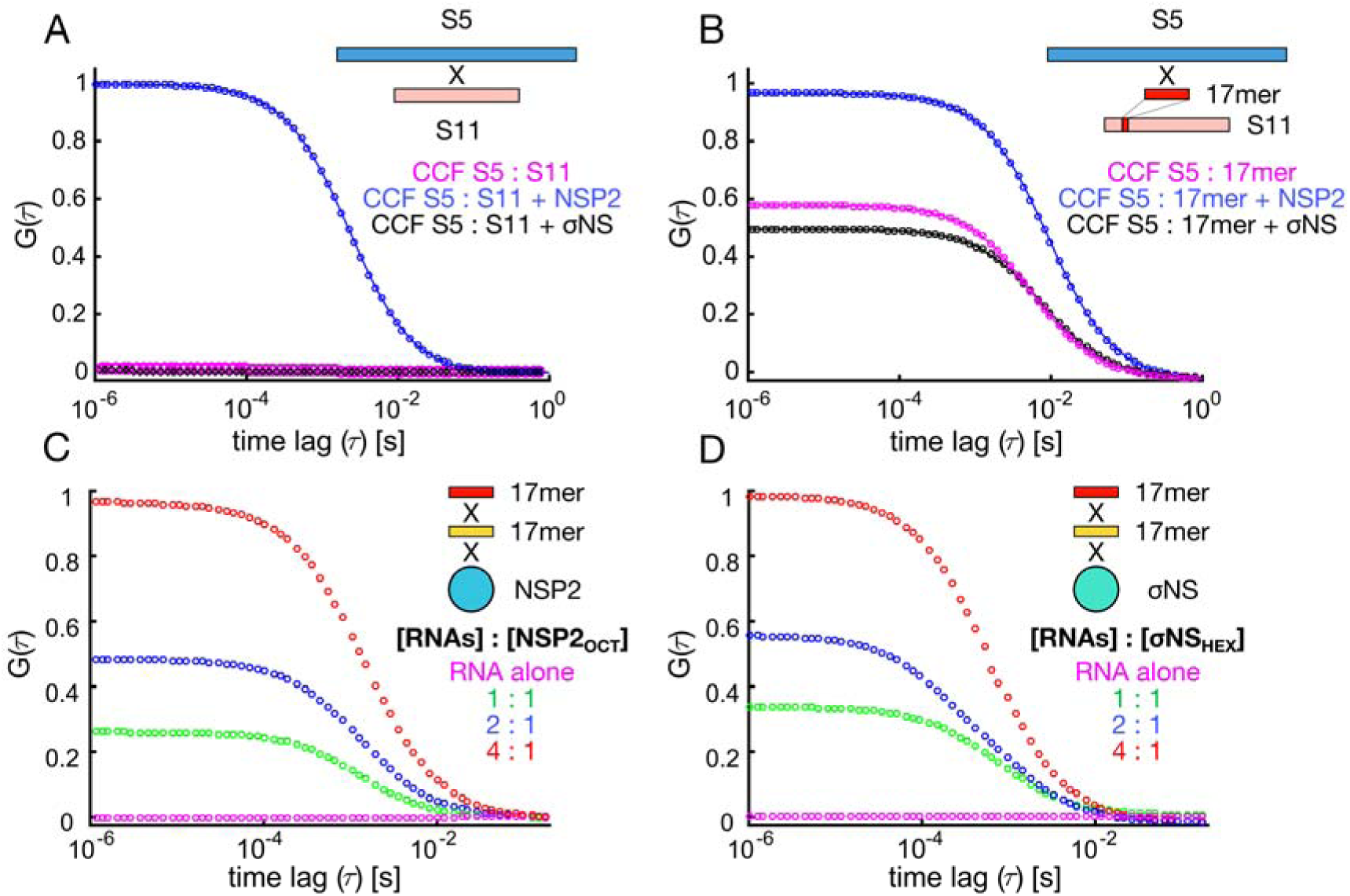
Probing RNA interactions mediated by RV NSP2 and ARV σNS. A: Inter-segment RNA-RNA interactions probed by fluorescence cross-correlation spectroscopy (FCCS). Normalised cross-correlation functions (CCF) are shown for interacting S5 & S11 RV RNAs. Equimolar mixtures of RV RNAs S5 and S11 (55 nM each) were incubated in the presence of 5 µM NSP2 (blue), or σNS (black), and diluted to achieve 1 nM RNA concentration. Under these conditions protein-free and σNS-bound S5 and S11 RNAs do not interact (magenta and black, respectively). B: Inter-molecular RNA interactions between a full-length S5 RNA and an unstructured 17-mer RNA derived from S11 RNA (Supporting Table S1). ssRNAs were incubated as described in (A), analysed by FCCS, yielding CCFs in the presence of NSP2 (blue), or σNS (black). C & D: Simultaneous binding of distinct 17-mer ssRNAs by NSP2 and σNS protein oligomers. Equimolar mixtures of Cy3-and Cy5-labelled non-complementary RNAs (Supporting Table S1) were incubated alone, and at variable RNA: protein oligomer ratios (hexameric σNS, σNS_HEX_ and octameric NSP2, NSP2_OCT_). CCF amplitudes were normalized by their respective ACFs, and the resulting amplitudes were then normalized to the highest CCF observed for 4:1 [RNA]: protein oligomer ratio, revealing co-diffusion of protein-bound distinct ssRNAs.

Given that σNS promotes strand-annealing of short RNA oligonucleotides (18), its failure to promote interactions between full-length genomic ssRNAs may be due to sequestration of complementary sequences within RNA secondary structure. We compared the strand-annealing activities of both proteins using shorter complementary RNA fragments derived from S5 (S5f, nucleotides 299 – 350) and S11 (S11f, nucleotides 31 – 77). Incubation of S5f and S11f (10 nM each) resulted in spontaneous hybridization (**Supplementary Figure S2**). To limit this, we investigated interactions between a full-length S5 and an unstructured S11-derived 17mer complementary to S5 (**Supporting Table S1**). NSP2 increases the amount of interacting S5: 17mer (**Figure 1B**, blue, respectively), while σNS was unable to promote this interaction beyond the level of spontaneous annealing (**Figure 1B**, black and magenta, respectively).

As RNA annealing activity typically involves simultaneous binding of two RNAs, we next examined whether NSP2 and σNS can bind multiple RNAs in solution. To distinguish strand-annealing from RNA binding, we designed differently-labelled non-complementary 17mer RNAs (**Supporting Table S1**). In the absence of protein, these RNAs did not interact (**Figure 1C & D**, magenta). Incubation of either NSP2 or σNS with an equimolar mixture of distinct RNAs at 2:1 to 4:1 protein oligomer: RNA ratio resulted in the largest fraction of co-diffusing oligomer-bound ssRNAs, indicating that both NSP2 and σNS can bind multiple unstructured RNAs. Having established that both proteins bind multiple RNAs with similarly high affinity, the observed failure of σNS to assist annealing of the 17mer to S5 ssRNA could be explained by its failure to remodel the target RNA sequence and increase its accessibility.

### σNS undergoes an RNA-induced hexamer-to-octamer transition

Having established that both NSP2 and σNS can bind multiple RNAs per oligomer, we further investigated σNS-RNP complex formation. NSP2 is a stable octamer in the presence or absence of RNA (11, 33, 42), while the σNS apoprotein is hexameric, although RNP stoichiometry is unknown (18). We incubated σNS hexamer with a stoichiometric excess of 20mer RNA (Materials and Methods) and analysed the RNP complexes using size-exclusion chromatography and dynamic light scattering. Surprisingly, σNS-RNP eluted earlier and had a greater hydrodynamic radius (*R* _h_) of ~ 8 nm than the apoprotein (*R* _h_ ~ 5 nm) (**Figure 2A**). This difference in *R*_h_ cannot be explained by the binding of multiple 20mer RNA molecules alone, suggesting that σNS undergoes a change in conformation or its oligomeric state.

**Figure. 2.**
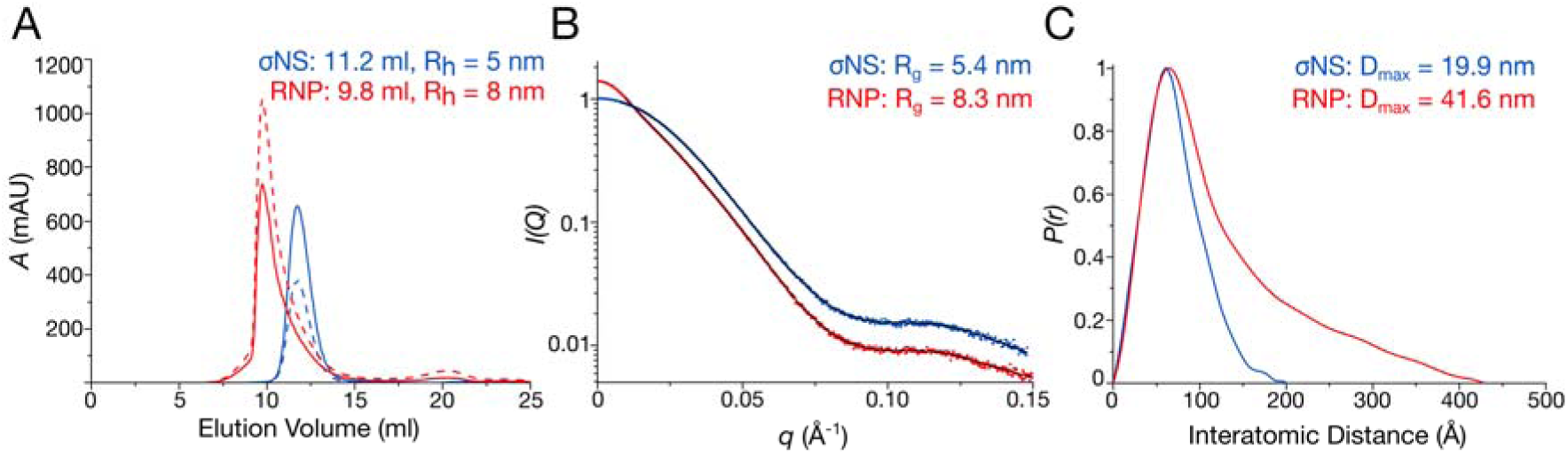
RNA binding results in assembly of a larger σNS oligomer. A: Size-exclusion chromatography elution profiles of σNS apoprotein (blue) and σNS ribonucleoprotein (RNP) complex (red). Absorbances at 260 nm and 280 nm are shown as dashed and continuous lines, respectively. Dynamic light scattering (DLS)-derived hydrodynamic radii (R_h_) are shown for each species. A second, minor peak corresponds to free, excess RNA. B: Small-angle X-ray scattering (SAXS) curves for σNS apoprotein (blue) and RNP complex (red), with respective fits shown in black. Scattering profiles are shown as the logarithm of the scattering intensity, *I*, as a function of the momentum transfer, q = 4πsin(θ)/λ. Radii of gyration (R_g_) values of both species are displayed. C: Normalised pair-wise distance distribution functions, P(r), calculated from the scattering curves of σNS apoprotein (blue) and RNP complex (red) showing an increase in maximum dimension (D_max_).

We then examined RNP complex formation by small-angle X-ray scattering (SAXS). Radii of gyration (*R* _g_) values for σNS apoprotein and RNP complex were 5.5 ± 0.03 nm and 7.6 ± 0.05 nm, respectively (**Figure 2B**). Guinier region analysis suggests that both σNS and σNS-RNP samples are monodisperse, confirming that the observed increase in size is not due to σNS aggregation when bound to RNA (**Supplementary Figure S3A**). Upon RNA binding, hexameric σNS further undergoes a 2-fold increase in its maximum distance (D_max_) value from 19.9 nm to 41.6 nm (**Figure 2C**). Further analysis of SAXS data indicates that both NSP2 and σNS are globular, however σNS-RNP complex formation results in a flexible, elongated particle (**Supplementary Figure S3B**). No further increase in size of the RNP was observed after incubation with a 40mer RNA (**Supplementary Figure S3C**). The assembled σNS-RNP complex appears to be more stable than σNS apoprotein, while NSP2 exhibited significantly decreased stability upon RNA binding (**Supplementary Figure S4**), explaining the severe aggregation of NSP2-RNP that precluded its characterisation by SAXS. Together, these data suggest that σNS undergoes an RNA-driven oligomerisation that is independent of the substrate RNA length.

To analyse the stoichiometries of σNS-RNP complexes, we used native electrospray ionization – ion mobility spectrometry – mass spectrometry (ESI-IMS-MS). A typical ESI-MS spectrum shows σNS hexamers along with additional oligomeric species, in agreement with previous analysis of σNS (18) (**Figure 3A & Supplementary Figure S5**). In contrast, the RNP contains a large population of octameric species with two RNAs bound (**Figure 3B**). A small fraction of hexameric species was bound to a single RNA. Together, IMS-MS and SAXS data suggest that σNS undergoes a hexamer-to-octamer transition upon RNP complex formation (**Figure 3**).

**Figure. 3.**
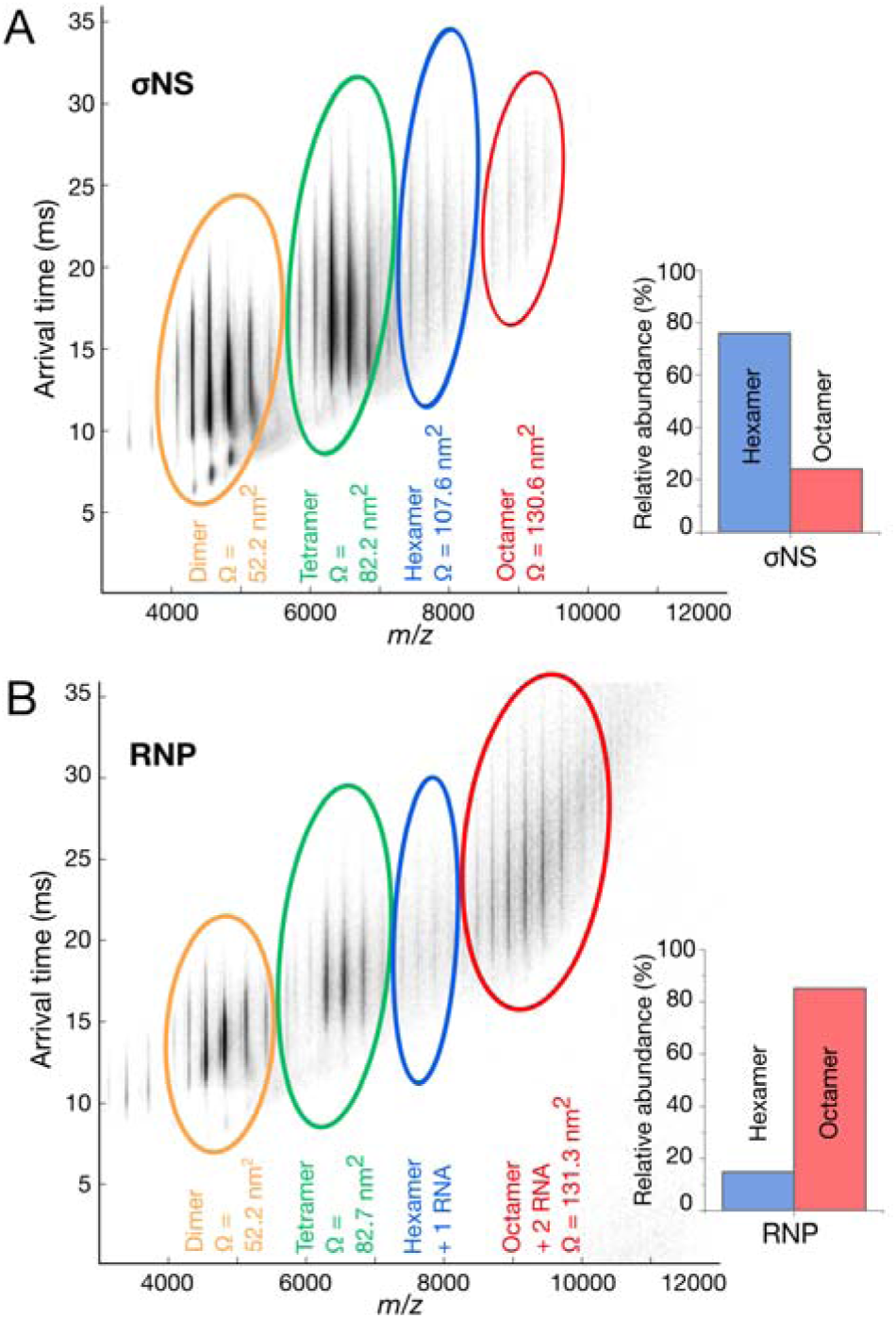
σNS RNP complex is predominantly octameric. A & B: Native electrospray ionisation (ESI) mass-spectra of σNS apoprotein (A) and RNP complex (B). Averaged collision cross-sections, CCS (Ω) are shown in nm^2^ for each species. Inset: Relative abundances of hexameric and octameric σNS oligomers. Smaller protein oligomers observed in both spectra are due to dissociation of higher order species during the ionization process.

We then used ion-mobility mass-spectrometry to estimate rotationally averaged collision cross-sections (CCSs) areas for each observed species (43). We compared CCSs derived from the ESI-IMS-MS spectra with the CCS values of σNS apoprotein and RNP complex calculated for SAXS-derived models, as described in Methods (**Figure 4A & B, Supplementary Figure S6**). The CCS values of the hexameric σNS (103.6 ± 2.8 nm^2^) are in good agreement with the CCS value estimated for the apoprotein SAXS envelope (102.2 ± 0.5 nm^2^) (**Figure 4C**). This is the same case for σNS-RNP, where the measured CCS value for octameric σNS bound in complex with 2 RNA molecules (120.5 ± 0.5 nm^2^) closely matches the size of the SAXS envelope generated for the RNP complex (119.4 ± 1.0 nm^2^). Thus, simultaneous binding of two RNAs results in the formation of stable, elongated octameric σNS-RNP complexes.

**Figure. 4.**
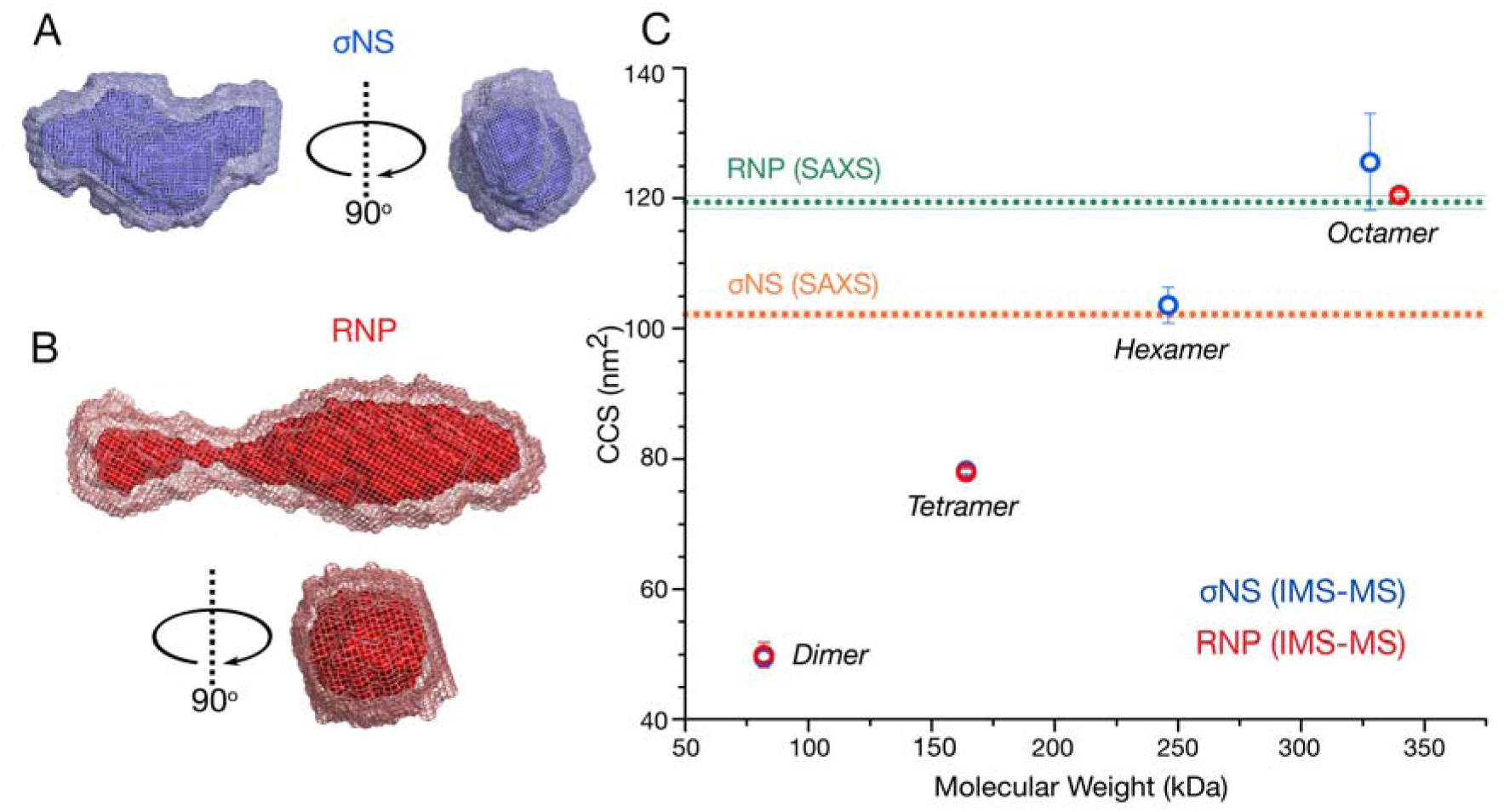
σNS undergoes a hexamer-to-octamer transition upon binding RNA. A & B: SAXS-derived *ab initio* models of hexameric σNS and σNS-RNP complex. 20 best models for each σNS apoprotein species (A) and σNS-RNP (B) were generated as described in Materials and Methods, averaged using DAMAVER (light mesh) and filtered using DAMFILT (superimposed dark surface). C: Collisional cross-sections (CCS) of σNS oligomers detected by ESI-IMS-MS. σNS apoprotein species are shown in blue and σNS-RNP are shown in red. Dashed horizontal lines denote CCS values estimated for the SAXS models of σNS apoprotein and the RNP complex, shown in (A) and (B), respectively. Masses of each oligomer and their charge states are summarised in Supporting Tables S4 & S5.

### Octameric σNS disrupts RNA structures more efficiently than its hexameric form

We then investigated the relationship between σNS oligomeric state and its helix-destabilizing activity. We designed a dual-labelled 36mer RNA stem-loop containing 3’-Atto532 and 5’Atto647N fluorophores (see Materials and Methods, **Figure 5A**), and incubated it with increasing amounts of σNS. Using electrophoretic mobility shift assays (EMSAs) to separate free RNAs from assembled RNP complexes (**Figure 5B & C**), we compared the helix-destabilizing activities of different σNS oligomers by estimating the FRET efficiencies of each band-shift (44, 45) (**Figure 5D** and **Supplementary Figure S7**). Titrations of σNS into this stem-loop produced three shifts that sequentially occurred at higher protein concentrations, corresponding to multiple oligomeric species. Given the presence of RNA-bound hexamers and octamers observed in the ESI-IMS-MS spectrum (**Figure 3**), we interpreted the first shift as hexameric σNS and the second shift as octameric σNS. The protein-free RNA had an apparent FRET efficiency (*E*_FRET(app)_) of 0.73 ± 0.03, suggesting that the stem-loop is folded. The RNA hexameric shift had *E*_FRET(app)_ of 0.40 ± 0.11, and this value decreased further for the octameric shift (*E*_FRET(app)_ = 0.28 ± 0.05), indicating that octameric σNS has a greater capacity for destabilizing RNA structure than hexameric σNS (**Figure 5D**).

**Figure. 5.**
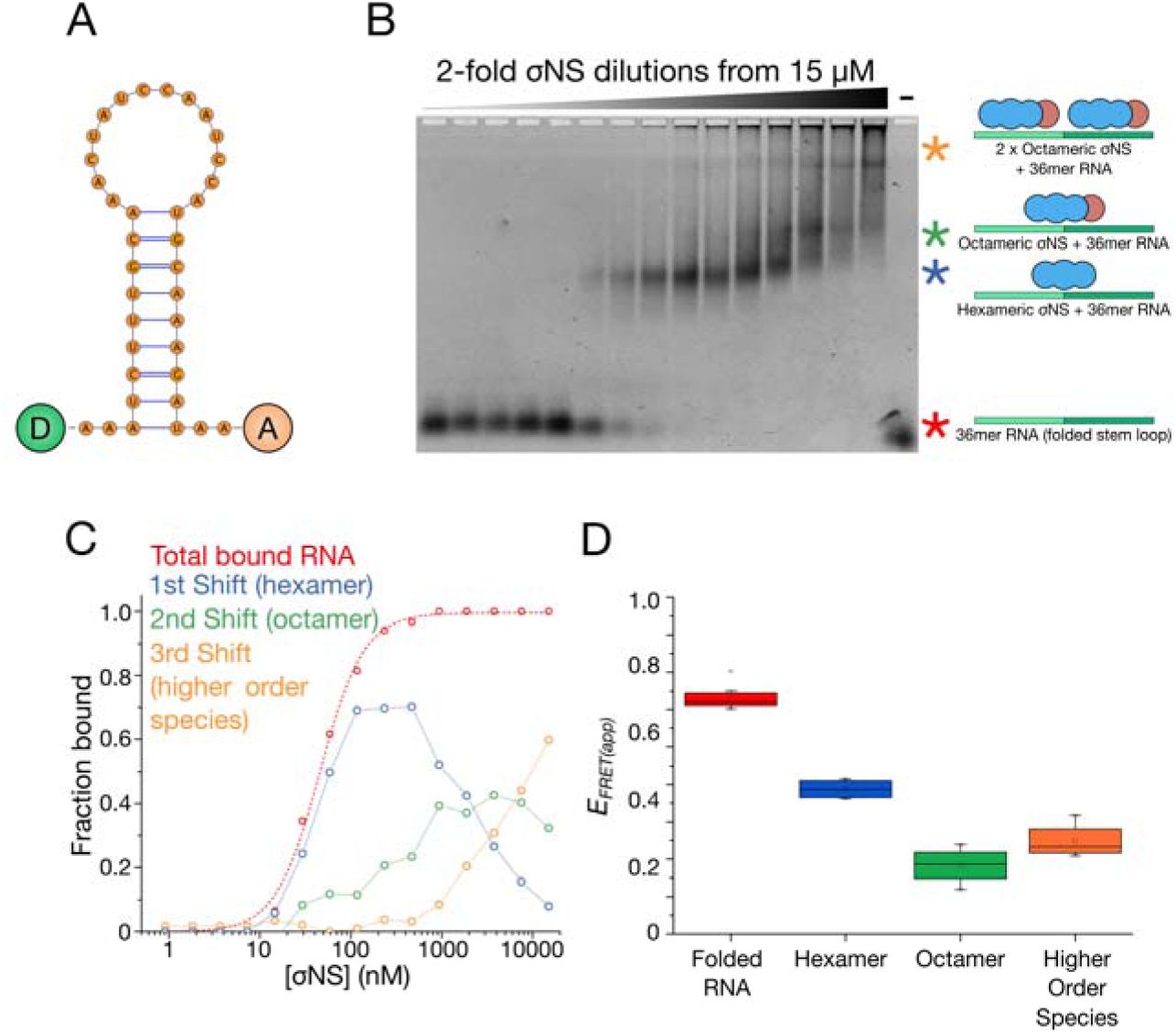
σNS oligomers have different helix-unwinding activities. A: Structure of the dual-labelled RNA stem-loop with 5′-donor (‘D’) and 3′-acceptor (‘A’) fluorophores, used for helix-unwinding assays monitored by Förster resonance energy transfer (FRET). B & C: Electrophoretic mobility shift assay (EMSA) of the dual-labelled stem-loop bound to σNS oligomers. Multiple shifts occur (blue, green and yellow asterisks) as σNS concentration increases. D: Helix-unwinding activities of different σNS oligomers formed at increasing σNS concentration. Apparent in-gel FRET efficiencies (*E_FRET(app)_*) of the unbound, hexamer-bound and octamer-bound RNA stem-loops were estimated for each band-shift shown in (B).

Given the RNA-binding footprint of σNS of ~20 nt (18, 46), a third shift was observed at elevated σNS concentrations (> 2 µM). This shift does not occur when σNS binds 20 nt RNA (**Supplementary Figure S8**), confirming that this shift is due to two σNS oligomers bound to the same 36 nt RNA. Similar protein saturation of 40mer RNA resulted in formation of a mixed population of higher-order species, as observed in SAXS (**Supplementary Figure S9**).

### NSP2 and σNS differ in modes of RNA unfolding

Having examined the helix-destabilising activity of σNS oligomers, we then investigated whether oligomer binding to a structured RNA is coupled to its unwinding activity. We used single-pair (sp)FRET to analyse the FRET states of discrete populations of protein-free and oligomer-bound RNAs using the dual-labelled RNA construct described above (**Figure 5A**). For stem-loop alone, a single high-FRET population is present (**Figure 6A**, grey histogram), in agreement with the MFE prediction that the RNA forms a stable hairpin with its 3’ and 5’ termini in proximity. In the presence of NSP2, two distinct FRET populations were observed – a high-FRET population similar to that of the stem-loop alone, and a low-FRET population (**Figure 6A**). Species-selective correlation analysis reveals that NSP2 is bound to RNA in both FRET populations (**Figure 6B**), indicating that NSP2 can bind to both folded and unfolded RNA stem-loops.

**Figure. 6.**
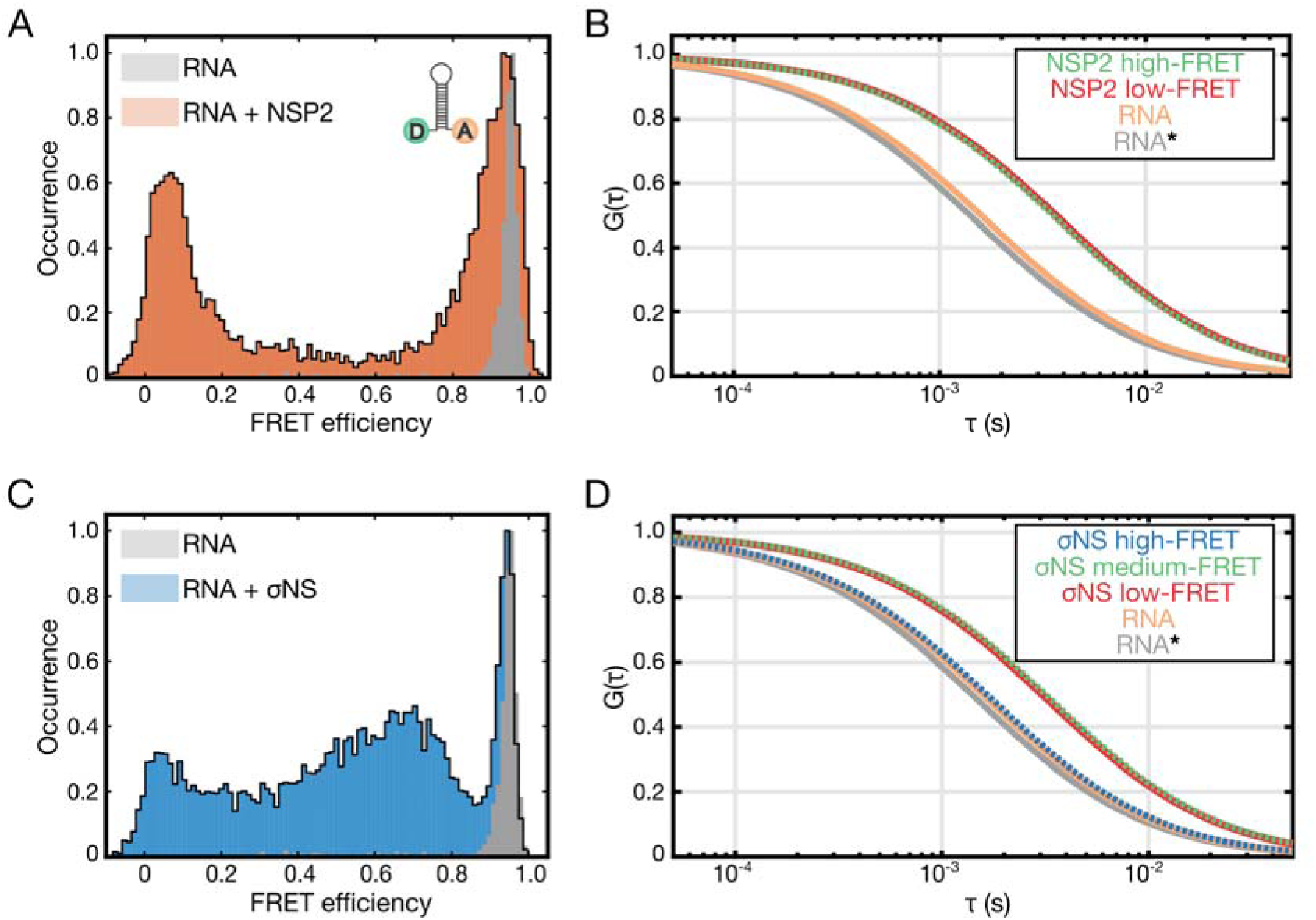
Helix-destabilising activities of σNS and NSP2, examined by single-pair FRET (spFRET) A: Histogram of spFRET efficiency of the dual-labelled RNA stem-loop (10 pM, shown in grey), and in the presence of 25 nM NSP2 (orange). B: Species-selective correlation analysis of the high-FRET (green autocorrelation function, ACF) and low-FRET (red ACF) populations, and freely diffusing RNA (orange). A typical ACF of a freely diffusing stem-loop is shown in grey (RNA*). Note rightward shift in ACFs of protein-bound stem-loops due to slower diffusion. C: Histogram of spFRET efficiency of the dual-labelled RNA stem-loop (see panel A), alone (grey) and in the presence of 25 nM σNS (blue). D: Species-selective correlation analysis of the high-FRET (blue ACF), intermediate FRET (green ACF) and low-FRET (red ACF) populations, and freely diffusing folded RNA (high-FRET, orange). A typical ACF of a freely diffusing stem-loop is shown in grey (RNA*). Only intermediate and low-FRET species are bound to σNS.

Similarly, we analysed RNA stem-loop destabilization by σNS (**Figure 6C**). While the high-FRET population persists in the presence of σNS, there are also a range of lower FRET populations, notably an intermediate-FRET and a low-FRET population. The intermediate-FRET population is dynamic on a sub-millisecond time scale, revealing an ensemble of partially unwound inter-converting σNS-bound RNA structures (**Supplementary Figure S10**). This is in agreement with previous results suggesting that even at higher protein concentrations (> 5 µM), σNS was unable to induce a single, unfolded RNA population (18). Species-selective correlation analysis indicates that σNS is bound to RNA in both the intermediate and low-FRET populations, but not the high-FRET (**Figure 6D**).

Ensemble FRET experiments conducted at a wider range of RNA and protein concentrations further demonstrate that NSP2 is more efficient at destabilizing RNA than σNS (**Supplementary Figure S11**). Collectively, these data suggest that NSP2 and σNS have different modes of RNA helix-destabilization. While NSP2 can bind to the folded stem-loop, its binding may not necessarily result in RNA unwinding. In contrast, σNS binding induces a range of unfolded and partially folded RNA conformations.

### NSP2 and σNS discriminate between RNA structures based on their relative stabilities

As RNA unfolding activities of both proteins require ssRNA binding, we investigated the relationship between RNA structure and binding affinity of NSP2 and σNS. We designed three fluorescently-labelled 20mer RNAs with different thermodynamic stabilities for use in binding measurements: an unstructured RNA, a metastable RNA (∆G= −3.8 kcal mol^-1^), and a stable hairpin structure (∆G= −8.1 kcal mol^-1^) (**Figure 7A**). All three RNAs bind NSP2 with near-identical affinities (*K*_D_ = 20 ± 3.2 nM, **Supporting Table S2**), indicating that NSP2 binds ssRNA independent of sequence (**Figure 7B**). σNS binds the unstructured and metastable RNAs with similar affinities (*K*_D_ = 37 ± 1.5 nM and 24 ± 2.6 nM, respectively, **Supporting Table S2**) but had lower affinity for the stable RNA (*K*_D_ = 137 ± 1.9 nM) (**Figure 7D**).

**Figure. 7.**
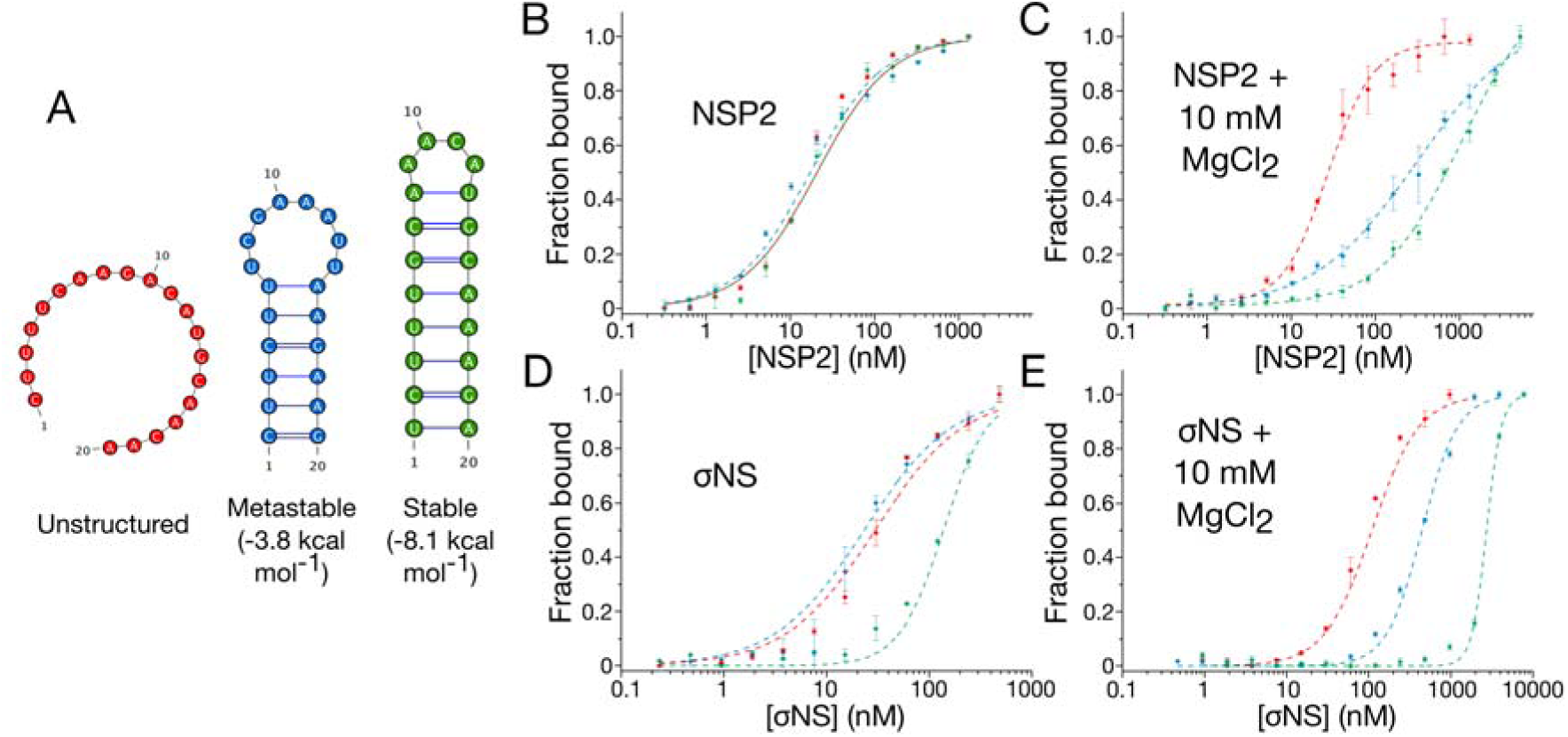
Stability of RNA structure determines preferential binding by NSP2 and σNS. A: Fluorescently-labelled unstructured (red), metastable (blue), and stable (green) 20-mer RNAs used for fluorescence anisotropy binding assays. B&D: NSP2 binds unstructured and stable RNAs with similar affinities. In contrast, stable secondary structure impedes σNS binding. C&E: Mg^2+^-dependent stabilisation of RNA structure impairs binding of ssRNAs by both NSP2 & σNS. Note the apparent affinity of both proteins for unstructured 20-mer remains largely unchanged upon addition of 10 mM MgCl_2_ Due to NSP2 aggregation at higher concentrations, protein titrations were only performed with [NSP2] up to 2 μM (panel C).

To further investigate RNA structural preferences of either protein, we then examined the affinities of NSP2 and σNS for these RNAs in the presence of Mg^2+^ ions, which stabilize RNA structures (47, 48). Although the *K*_D_ values of both NSP2 and σNS for unstructured RNA remains largely unchanged (1.4-and 1.5-fold increase, respectively), there is a 10-to 30-fold decrease in affinity of both NSP2 and σNS for the metastable and stable RNAs (**Figure 7C & 7E** & **Supporting Table S2**). This suggests that although both NSP2 and σNS exhibit preferential binding to unstructured RNAs, NSP2 can bind stable hairpins better than σNS. This reduction in affinity cannot be explained by Mg^2+^-induced dissociation of protein oligomers (33), as binding affinities of either protein for unstructured RNA remain largely unaffected in the presence of Mg^2+^.

Previous structural studies of NSP2 have demonstrated that sequence-independent ssRNA-binding occurs via a positively-charged groove (11, 42). Although there is no such structural information for σNS, NSP2 and σNS both bind ssRNA with high affinity and without apparent sequence specificity, potentially via multiple electrostatic contacts. We therefore examined the dependency of binding affinity on ionic strength by measuring the dissociation constant of NSP2 and σNS for an unstructured 20mer ssRNA molecule at different ionic strength (**Figure 8**). We observed the expected relationships between binding affinity and ionic strength (Eq., indicating that NSP2 binding involves at most two salt bridges, whereas σNS binds RNA via 3 – 4 ion contacts (**Figure 8C**) (32). Under physiological ionic strength, NSP2 binds ssRNA with apparent free energy of binding, ∆G= −9.52 kcal mol^-1^, of which there is an estimated ~17.5% electrostatic component **(Supporting Table S3**). Under the same conditions, σNS binds RNA with ∆G= −9.50 kcal mol^-1^, of which there is a ~30.4% electrostatic contribution (**Supporting Table S3**). This indicates that despite their near-identical affinities for RNA, σNS has a greater electrostatic contribution to the overall free energy of binding relative to that of NSP2. Together, these results suggest that NSP2 and σNS interact with RNAs differently, allowing them to discriminate between ssRNAs based on their propensities to form stable secondary structures (**Figure 9**).

**Figure. 8.**
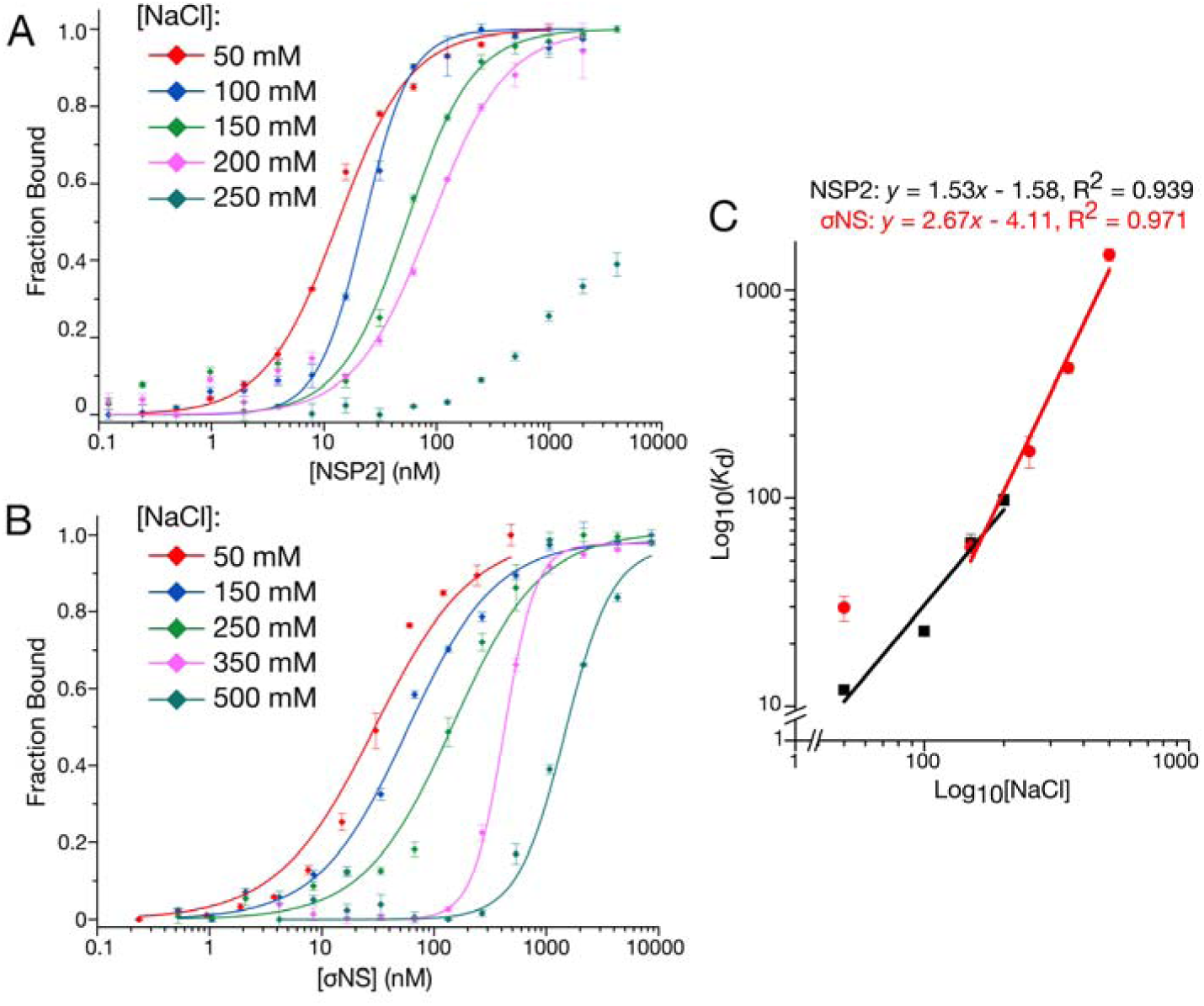
NSP2 and σNS display different electrostatic contributions to RNA binding. A & B: Salt-dependence of NSP2 (A) and σNS (B) binding to unstructured 20-mer ssRNA measured by fluorescence anisotropy. C: Linear correlation between log(*K*_D_) and log([NaCl]) for both NSP2 (black) and σNS (red). Derived mean K_d_ ± sd (N = 3) values are summarised in Supporting Table S3. The number of salt bridges contributing to RNA binding is estimated from fitted slopes, corresponding to <2 for NSP2 and 3 –4 for σNS.

## Discussion

Inter-molecular RNA-RNA interactions have been postulated to underpin the selection and assembly of multi-segmented genomes in viruses comprising the *Reoviridae* family (5, 17, 49, 50). Recent studies on rotaviruses suggest that NSP2 promotes RNA-RNA interactions between its full-length genomic ssRNA segment precursors (17). Both NSP2 and σNS facilitate conformational rearrangements of ssRNAs *in vitro* (17, 18) and mediate formation of inter-molecular RNA contacts between short, synthetic RNA fragments, including those derived from ARV genome and containing complementary sequences that may be involved in assembly of multiple RNAs (18). However, such interacting sequences have not been yet identified in mammalian or avian reoviruses, and have only recently been mapped for a number of inter-segment contacts within the rotavirus genome (17). Therefore we employed these RV RNAs as a model to directly compare the abilities of rotavirus NSP2 and avian reovirus σNS to mediate RNA-RNA interactions within the context of full-length RNA genomic precursors.

Here we have shown that despite the ostensibly similar RNA chaperone-like activities of NSP2 and σNS, only NSP2 is capable of promoting inter-segment interactions between RV RNAs. To gain insights into the mechanisms underpinning this selective, protein-mediated strand-annealing reaction, we performed a side-by-side comparison of the RNA binding, helix-destabilizing and strand-annealing activities of both proteins.

Previous studies conducted with ssRNA substrates suggest that both NSP2 and σNS bind ssRNA without apparent sequence specificity (11, 12, 18, 33, 42, 46). Using RNA substrates with different secondary structure stabilities, here we have demonstrated that σNS displays reduced affinity towards highly stable hairpins, whereas NSP2 binds ssRNAs regardless of their propensity to form secondary structure. Further stabilization of RNA structure with Mg^2+^ ions reveals that both proteins indeed preferentially bind unstructured RNAs. The observed change in the affinity is consistent with previously reported inhibition of the strand displacement activity of σNS in the presence of MgCl_2_ (42). NSP2 was shown to dissociate into smaller oligomers in the presence of magnesium (33). However, NSP2 binding to unstructured 20-mer RNA in the presence of 10 mM MgCl_2_ remains largely unaffected (Figure 7C & 7E), strongly suggesting that the observed inhibitory effects of Mg^2+^ ions are due to RNA secondary structure stabilization. These structural preferences of NSP2 and σNS are consistent with the observed helix-destabilization upon protein binding to a stable RNA stem-loop. Incubation of the stem-loop with large molar excess of either protein results in the RNA helix disruption (**Supplementary Figure S11**). However, spFRET and FCS analyses reveal that NSP2 can efficiently bind both folded (high FRET) and unfolded (low FRET) RNA stem-loops. The initial NSP2 binding may not necessarily result in substantial RNA unwinding (**Figure 6**), which is ultimately achieved at higher protein concentrations (**Supplementary Figure S11**). In contrast, σNS binding results in gradual unfolding, producing a mixed population of partially unfolded RNA intermediates, failing to completely unfold RNA even at high molar excess, further indicating that both proteins have different modes of helix disruption (**Figure 6**).

Given that helix-destabilizing activity of both proteins is likely to be coupled to ssRNA binding, we examined how these proteins interact with unstructured RNAs. While NSP2 interacts with two RNAs as an octamer (33), σNS hexamer only binds a single RNA and undergoes a hexamer-to-octamer transition that appears to be a prerequisite for binding a second RNA molecule. Octamer formation is concomitant with the increased helix-destabilizing activity, potentially providing additional RNA-binding surface (51, 52), hence increased capacity to compete with RNA secondary structure formation (**Figure 9**). Thus, failure of σNS to promote a specific inter-segment duplex formation can be attributed to its reduced capacity to interact with and disrupt stable intramolecular RNA structures.

**Figure. 9.**
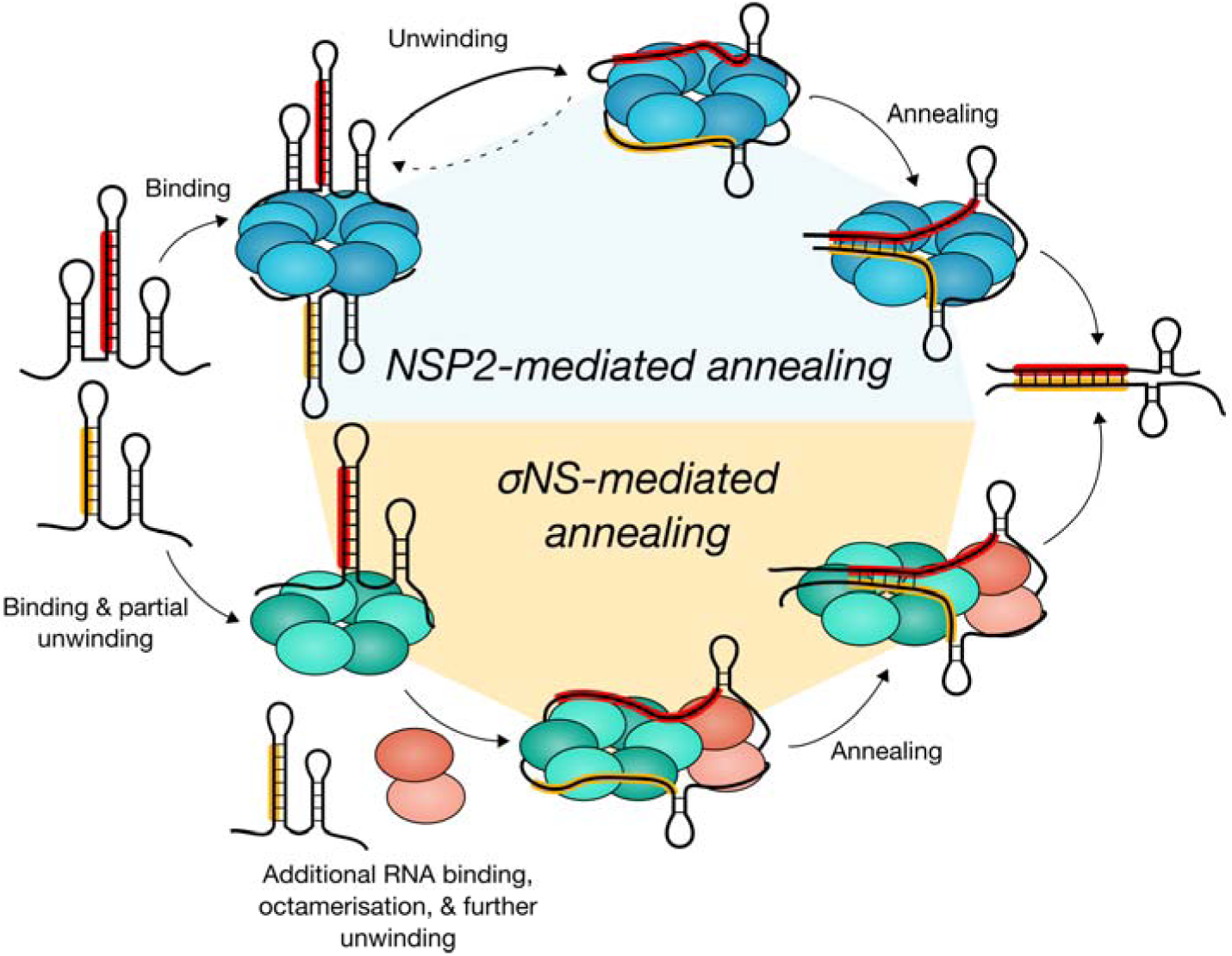
NSP2 and σNS employ different mechanisms to promote RNA-RNA interactions. NSP2 (blue) and σNS (green) can bind multiple RNAs per oligomer. NSP2 octamer binding results in efficient RNA unwinding, thereby promoting duplex formation between complementary sequences (highlighted in red and yellow) within interacting genomic segment ssRNAs. In contrast, efficient RNA unwinding by σNS requires a hexamer-to-octamer transition triggered by additional RNA binding. Failure of σNS oligomers to fully disrupt complementary sequences sequestered within RNA secondary structure results in abrogation of strand-annealing activity.

### RNA secondary structure stability modulates chaperone activity

Given the mechanistic differences between NSP2 and σNS, it appears that a major determinant of efficient protein-mediated inter-segment annealing is RNA structural stability. Intramolecular RNA structure can regulate binding by cognate RNA chaperone proteins, therefore affecting refolding and inter-molecular duplex annealing. Thus, RNA secondary structure stability together with chaperone binding mode may serve to fine-tune the matchmaking activities of RNA chaperones that would otherwise interact with RNA without sequence preference (53, 54). This may regulate specific RNA-RNA interactions required for a highly accurate assembly of a complete segmented RNA genome. Other RNA chaperones, including *E. coli* StpA and Moloney murine leukemia virus NC, both of which bind preferentially to unstructured regions of RNA (55, 56), exhibit similar relationships between chaperone activity and RNA structural stability (57, 58). This suggests that fine-tuning of RNA remodelling by its stability may be a general feature of RNA chaperone activity.

### RNA-driven oligomerisation

Upon binding of two ssRNAs hexameric σNS assembles into octameric RNP complexes with defined stoichiometry. This mechanism of RNA-driven oligomerisation is distinct from that of other viral RNA-binding proteins that assemble into large, non-discrete, higher-order oligomers in the presence of RNA (59–62). We propose that σNS hexamer assembles as a trimer of dimers (18), in dynamic equilibrium with a low octameric population (**Figure 3**). RNA binding results in equilibrium shift towards octameric species, which appear to be stabilized by RNP complex formation. This represents a novel mechanism of modulating RNA chaperone activity, whereby the oligomeric state of the protein defines its unwinding efficiency. Similar to ARV σNS, mammalian reovirus σNS and bluetongue virus NS2 have also been reported to exist in a range of oligomeric states (63–66). It is possible that different σNS oligomers may play distinct roles during viral replication.

Other functionally analogous proteins encoded by different members of the *Reoviridae* family form octamers, similar to NSP2 (notably P9–1 and Pns9 proteins (67, 69)). Despite their similar toroidal architecture, NSP2 presents a continuous, basic RNA binding groove on the surface, while P9–1 binds ssRNA within a positively charged inner pore. Analysis of RNA binding by both NSP2 and σNS also reveals different number of salt bridges involved in protein-RNA interactions (**Figure 8**), with each protein predominantly interacting non-electrostatically. Such sequence-independent, non-electrostatic interactions with ribose and base moieties may explain the preference of both NSP2 and σNS for unfolded ssRNA over dsRNA substrates that are only accessible via the A-form backbone. Indeed, Raman difference analysis of σNS-bound RNA (**Supplementary Figure S12**) reveals decrease in the band intensities corresponding to A-form backbone vibrations, with many base vibrations being affected, and only small changes in the bands arising from phosphate vibrations.

Our results suggest that different members of the *Reoviridae* family may exploit distinct mechanisms of regulating RNA chaperone activities underpinning genome segment assortment. We propose that the stability of RNA structure together with distinct unwinding mechanisms underpins the observed selectivity of RNA-RNA interactions. This may serve to regulate selection of genomic RNA segments to achieve assembly of a complete set of cognate genomic RNAs with high fidelity.

## DATA ACCESS

SAXS models reported in this paper can be accessed at SASBDB (https://www.sasbdb.org/), ID numbers: SASDDT5 (σNS-RNP), SASDDU5 (σNS apoprotein).

## ACKNOWLEDGEMENTS

We thank Nikul Khunti (B21 SAXS beamline support scientists team, Diamond Light Source, UK) for technical assistance with SAXS data acquisition.

## Author contributions

J.P.K.B. and A.Borodavka designed and carried out experiments, analyzed data and jointly wrote the manuscript. A.Barth and D.C.L. contributed novel analytical tools and A.Barth analyzed spFRET data. A.C. contributed novel analytical tools and analyzed ESI-IMS-MS data. A.Borodavka and R.T. managed the project. P.M. collected and analyzed Raman spectroscopy data. All authors contributed ideas, discussed the results and were involved in writing of the manuscript.

## References

1. Desselberger, U. (2014) Rotaviruses. Virus Res., 190, 75–96.

2. Mertens, P. (2004) The dsRNA viruses. Virus Res., 101, 3–13.

3. McDonald, S.M., Nelson, M.I., Turner, P.E. and Patton, J.T. (2016) Reassortment in segmented RNA viruses: mechanisms and outcomes. Nat. Rev. Microbiol., 14, 448–460.

4. McDonald, S.M. and Patton, J.T. (2011) Assortment and packaging of the segmented rotavirus genome. Trends Microbiol., 19, 136–144.

5. Sung, P.Y. and Roy, P. (2014) Sequential packaging of RNA genomic segments during the assembly of bluetongue virus. Nucleic Acids Res., 42, 13824–13838.

6. Lourenco, S. and Roy, P. (2011) In vitro reconstitution of Bluetongue virus infectious cores. Proc. Natl. Acad. Sci. U. S. A., 108, 13746–51.

7. Anzola, J. V, Xu, Z.K., Asamizu, T. and Nuss, D.L. (1987) Segment-specific inverted repeats found adjacent to conserved terminal sequences in wound tumor virus genome and defective interfering RNAs. Proc Natl Acad Sci U S A, 84, 8301–8305.

8. Tourís-Otero, F., Cortez-San Martín, M., Martínez-Costas, J. and Benavente, J. (2004) Avian reovirus morphogenesis occurs within viral factories and begins with the selective recruitment of σNS and λA to µNS inclusions. J. Mol. Biol., 341, 361–374.

9. Netherton, C.L. and Wileman, T. (2011) Virus factories, double membrane vesicles and viroplasm generated in animal cells. Curr. Opin. Virol., 1, 381–387.

10. Silvestri, L.S., Taraporewala, Z.F. and Patton, J.T. (2004) Rotavirus Replication: Plus-Sense Templates for Double-Stranded RNA Synthesis Are Made in Viroplasms. J. Virol., 78, 7763–7774.

11. Jiang, X., Jayaram, H., Kumar, M., Ludtke, S.J., Estes, M.K. and Venkataram Prasad, B. V (2006) Cryoelectron Microscopy Structures of Rotavirus NSP2-NSP5 and NSP2-RNA Complexes: Implications for Genome Replication. J. Virol., 80, 10829–10835.

12. Taraporewala, Z.F., Jiang, X., Vasquez-Del Carpio, R., Jayaram, H., Prasad, B.V.V. and Patton, J.T. (2006) Structure-function analysis of rotavirus NSP2 octamer by using a novel complementation system. J. Virol., 80, 7984–7994.

13. Touris-Otero, F., Martínez-Costas, J., Vakharia, V.N. and Benavente, J. (2004) Avian reovirus nonstructural protein μNS forms viroplasm-like inclusions and recruits protein σNS to these structures. Virology, 319, 94–106.

14. Miller, C.L., Broering, T.J., Parker, J.S.L., Arnold, M.M. and Nibert, M.L. (2003) Reovirus σNS protein localizes to inclusions through an association requiring the μNS amino terminus. J. Virol., 77, 4566–76.

15. Desmet, E.A., Anguish, L.J. and Parker, J.S.L. (2014) Virus-mediated compartmentalization of the host translational machinery. MBio, 5, 1–11.

16. Miller, C.L., Arnold, M.M., Broering, T.J., Hastings, C.E. and Nibert, M.L. (2010) Localization of mammalian orthoreovirus proteins to cytoplasmic factory-like structures via nonoverlapping regions of microNS. J. Virol., 84, 867–882.

17. Borodavka, A., Dykeman, E.C., Schrimpf, W. and Lamb, D.C. (2017) Protein-mediated RNA folding governs sequence-specific interactions between rotavirus genome segments. Elife, 6, 1–22.

18. Borodavka, A., Ault, J., Stockley, P.G. and Tuma, R (2015) Evidence that avian reovirus σNS is an RNA chaperone: implications for genome segment assortment. Nucleic Acids Res., 43, 7044–7057.

19. Borodavka, A., Singaram, S.W., Stockley, P.G., Gelbart, W.M., Ben-Shaul, A. and Tuma, R. (2016) Sizes of Long RNA Molecules Are Determined by the Branching Patterns of Their Secondary Structures. Biophys. J., 111, 2077–2085.

20. Konarev, P. V, Volkov, V. V, Sokolova, A. V, Koch, M.H.J. and Svergun, D.I. (2003)PRIMUSI⍰: a Windows PC-based system for small-angle scattering data analysis. J. Appl. Crystallogr., 36, 1277–1282.

21. Konarev, P. V., Petoukhov, M. V., Volkov, V. V. and Svergun, D.I. (2006) ATSAS 2.1, a program package for small-angle scattering data analysis. J. Appl. Crystallogr., 39, 277–286.

22. Petoukhov, M. V., Franke, D., Shkumatov, A. V., Tria, G., Kikhney, A.G., Gajda, M., Gorba, C., Mertens, H.D.T., Konarev, P. V. and Svergun, D.I. (2012) New developments in the ATSAS program package for small-angle scattering data analysis. J. Appl. Crystallogr., 45, 342–350.

23. Franke, D. and Svergun, D.I. (2009) DAMMIF, a program for rapid ab-initio shape determination in small-angle scattering. J. Appl. Crystallogr., 42, 342–346.

24. Volkov, V. V., and Svergun, D.I. (2003) Uniqueness of ab initio shape determination in small-angle scattering. J. Appl. Crystallogr., 36, 860–864.

25. Morgner, N. and Robinson, C. V. (2012) Massign: An assignment strategy for maximizing information from the mass spectra of heterogeneous protein assemblies. Anal. Chem., 84, 2939–2948.

26. Ruotolo, B.T., Benesch, J.L.P., Sandercock, A.M., Hyung, S.-J. and Robinson, C. V (2008) Ion mobility-mass spectrometry analysis of large protein complexes. Nat. Protoc., 3, 1139–1152.

27. Bush, M.F., Hall, Z., Giles, K., Hoyes, J., Robinson, C. V. and Ruotolo, B.T. (2010) Collision cross sections of proteins and their complexes: A calibration framework and database for gas-phase structural biology. Anal. Chem., 82, 9557–9565.

28. Smith, D., Knapman, T., Campuzano, I., Malham, R., Berryman, J., Radford, S.E. and Ashcroft, A. (2009) Deciphering drift time measurements from travelling wave ion mobility spectrometry-mass spectrometry studies. Eur. J. Mass Spectrom., 15, 113.

29. Ruotolo, B.T. and Robinson, C. V (2006) Aspects of native proteins are retained in vacuum. Curr. Opin. Chem. Biol., 10, 402–408.

30. Marklund, E.G., Degiacomi, M.T., Robinson, C. V., Baldwin, A.J. and Benesch, J.L.P. (2015) Collision cross sections for structural proteomics. Structure, 23, 791–799.

31. Davidovich, C., Zheng, L., Goodrich, K.J. and Cech, T.R. (2013) Promiscuous RNA binding by Polycomb repressive complex 2. Nat. Struct. Mol. Biol., 20, 1250–1257.

32. Record, M.T., Lohman, T.M. and Haseth, P. de (1976) Ion effects on ligand-nucleic acid interactions. J. Mol. Biol., 107, 145–158.

33. Schuck, P., Taraporewala, Z., McPhie, P. and Patton, J.T. (2001) Rotavirus Nonstructural Protein NSP2 Self-assembles into Octamers that Undergo Ligand-induced Conformational Changes. J. Biol. Chem., 276, 9679–9687.

34. Kudryavtsev, V., Sikor, M., Kalinin, S., Mokranjac, D., Seidel, C.A.M. and Lamb, D.C. (2012) Combining MFD and PIE for accurate single-pair förster resonance energy transfer measurements. ChemPhysChem, 13, 1060–1078.

35. Voith von Voithenberg, L., Sánchez-Rico, C., Kang, H.-S., Madl, T., Zanier, K., Barth, A., Warner, L.R., Sattler, M. and Lamb, D.C. (2016) Recognition of the 3⍰ splice site RNA by the U2AF heterodimer involves a dynamic population shift. Proc. Natl. Acad. Sci., 113, E7169–E7175.

36. Nir, E., Michalet, X., Hamadani, K., Laurence, T.A., Neuhauser, D., Kovchegov, Y. and Weiss, S. (2006) Shot-Noise Limited Single-Molecule FRET HIstograms: Comparison between Theory and Experiments. J. Phys. Chem. B, 110, 22103–22124.

37. Tomov, T.E., Tsukanov, R., Masoud, R., Liber, M., Plavner, N. and Nir, E. (2012) Disentangling subpopulations in single-molecule FRET and ALEX experiments with photon distribution analysis. Biophys. J., 102, 1163–1173.

38. Eggeling, C., Fries, J.R., Brand, L., Gunther, R. and Seidel, C.A.M. (1998) Monitoring conformational dynamics of a single molecule by selective fluorescence spectroscopy. Proc. Natl. Acad. Sci. U. S. A., 95, 1556–1561.

39. Laurence, T.A., Kwon, Y., Yin, E., Hollars, C.W., Camarero, J.A. and Barsky, D. (2007) Correlation spectroscopy of minor fluorescent species: Signal purification and distribution analysis. Biophys. J., 92, 2184–2198.

40. Schrimpf, Waldemar (2018) PAM: A Framework for Integrated Analysis of Imaging, Single-Molecule, and Ensemble Fluorescence Data. Biophys. J.

41. Palacký, J., Mojzeš, P. and Bok, J. (2011) SVD-based method for intensity normalization, background correction and solvent subtraction in Raman spectroscopy exploiting the properties of water stretching vibrations. J. Raman Spectrosc., 42, 1528–1539.

42. Jayaram, H., Taraporewala, Z., Patton, J.T. and Prasad, B.V.V. (2002) Rotavirus protein involved in genome replication and packaging exhibits a HIT-like fold. Nature, 417, 311–5.

43. Schiffrin, B., Calabrese, A.N., Devine, P.W.A., Harris, S.A., Ashcroft, A.E., Brockwell, D.J. and Radford, S.E. (2016) Skp is a multivalent chaperone of outer-membrane proteins. Nat. Struct. Mol. Biol., 23, 786–793.

44. Ghetu, A.F., Arthur, D.C., Kerppola, T.K. and Glover, J.N.M. (2002) Probing FinO-FinP RNA interactions by site-directed protein-RNA crosslinking and in-gel FRET. RNA, 8, 816–823.

45. Radman-Livaja, M., Biswas, T., Mierke, D. and Landy, A. (2005) Architecture of recombination intermediates visualized by in-gel FRET of λ integrase-Holliday junction-arm DNA complexes. Proc. Natl. Acad. Sci. U. S. A., 102, 3913–3920.

46. Gillian, A.L., Schmaechel, S.C., Livny, J., Schiff, L.A. and Nibert, M.L. (2000) Reovirus protein σNS binds in multiple copies to single stranded RNA and shares properties with single stranded DNA binding proteins. J. Virol., 74, 5939–5948.

47. Draper, D.E. (2004) A guide to ions and RNA structure. RNA, 10, 335–343.

48. Misra, V.K. and Draper, D.E. (1998) On the Role of Magnesium Ions in RNA Stability. Biopolymers, 48, 113–135.

49. Fajardo, T., Sung, P.-Y. and Roy, P. (2015) Disruption of Specific RNA-RNA Interactions in a Double-Stranded RNA Virus Inhibits Genome Packaging and Virus Infectivity. PLOS Pathog., 11, e1005321.

50. Fajardo, T.J., Al Shaikhahmed, K. and Roy, P. (2016) Generation of infectious RNA complexes in Orbiviruses: RNA-RNA interactions of genomic segments. Oncotarget, 7, 72559–72570

51. Russell, R. (2008) RNA misfolding and the action of chaperones. Front. Biosci., 13, 1–20.

52. Rajkowitsch, L., Chen, D., Stampfl, S., Semrad, K., Waldsich, C., Mayer, O., Jantsch, M.F., Konrat, R., Bläsi, U. and Schroeder, R. (2007) RNA chaperones, RNA annealers and RNA helicases. RNA Biol., 4, 118–30.

53. Müller, U.F. and Göringer, H.U. (2002) Mechanism of the gBP21-mediated RNA / RNA annealing reaction⍰: matchmaking and charge reduction. Nucleic Acids Res., 30, 447–455.

54. Peng, Yi, Curtis, J.E., Fang, X. and Woodson, S.A. (2014) Structural model of an mRNA in complex with the bacterial chaperone Hfq. Proc Natl Acad Sci U S A, 111, 17134–9.

55. Mayer, O., Rajkowitsch, L., Lorenz, C., Konrat, R. and Schroeder, R. (2007) RNA chaperone activity and RNA-binding properties of the E. coli protein StpA. Nucleic Acids Res., 35, 1257–1269.

56. D’Souza, V. and Summers, M.F. (2004) Structural basis for packaging the dimeric genome of Moloney murine leukaemia virus. Nature, 431, 586–590.

57. Grossberger, R., Mayer, O., Waldsich, C., Semrad, K., Urschitz, S. and Schroeder, R. (2005) Influence of RNA structural stability on the RNA chaperone activity of the Escherichia coli protein StpA. Nucleic Acids Res., 33, 2280–2289.

58. Grohman, J.K., Gorelick, R.J., Lickwar, C.R., Lieb, J.D., Bower, B.D., Znosko, B.M. and Weeks, K.M. (2013) A guanosine-centric mechanism for RNA chaperone function. Science, 340, 190–5.

59. Milles, S., Jensen, M.R., Communie, G., Maurin, D., Schoehn, G., Ruigrok, R.W.H. and Blackledge, M. (2016) Self-Assembly of Measles Virus Nucleocapsid-like Particles: Kinetics and RNA Sequence-Dependence. Angew. Chemie Int. Ed., 55, 9356–9360.

60. Reguera, J., Cusack, S. and Kolakofsky, D. (2014) Segmented negative strand RNA virus nucleoprotein structure. Curr. Opin. Virol., 5, 7–15.

61. Durham, A.C., Finch, J.T. and Klug, A. (1971) States of aggregation of tobacco mosaic virus protein. Nat. New Biol., 229, 37–42.

62. Butler, P.J.G. and Klug, A. (1971) Assembly of the particle of tobacco mosaic virus from RNA and disks of protein. Nat. New Biol., 229, 47–50.

63. Mumtsidu, E., Makhov, A.M., Roessle, M., Bathke, A. and Tucker, P.A. (2007) Structural features of the Bluetongue virus NS2 protein. J. Struct. Biol., 160, 157–167.

64. Butan, C. and Tucker, P. (2010) Insights into the role of the non-structural protein 2 (NS2) in Bluetongue virus morphogenesis. Virus Res., 151, 109–117.

65. Butan, C., Van Der Zandt, H. and Tucker, P. a (2004) Structure and assembly of the RNA binding domain of bluetongue virus non-structural protein 2. J. Biol. Chem., 279, 37613–37621.

66. Gillian, A.L. and Nibert, M.L. (1998) Amino terminus of reovirus nonstructural protein sigma NS is important for ssRNA binding and nucleoprotein complex formation. Virology, 240, 1–11.

67. Akita, F., Higashiura, A., Shimizu, T., Pu, Y., Suzuki, M., Uehara-ichiki, T., Sasaya, T., Kanamaru, S., Arisaka, F., Tsukihara, T., et al. (2012) Crystallographic Analysis Reveals Octamerization of Viroplasm Matrix Protein P9-1 of Rice BlacKStreaked Dwarf Virus. J. Virol., 86, 746–756.

68. Akita, F., Miyazaki, N., Hibino, H., Shimizu, T., Higashiura, A., Uehara-Ichiki, T., Sasaya, T., Tsukihara, T., Nakagawa, A., Iwasaki, K., et al. (2011) Viroplasm matrix protein Pns9 from rice gall dwarf virus forms an octameric cylindrical structure. J. Gen. Virol., 92, 2214–2221.

69. Wu, J., Li, J., Mao, X., Wang, W., Cheng, Z., Zhou, Y. and Zhou, X. (2013) Viroplasm Protein P9-1 of Rice Black-Streaked Dwarf Virus Preferentially Binds to Single-Stranded RNA in Its Octamer Form, and the Central Interior Structure Formed by This Octamer Constitutes the Major RNA Binding Site. J.Virol., 87, 12885–12899.

